# Foraging task specialization in honey bees (*Apis mellifera*): the contribution of floral rewards on the learning performance of pollen and nectar foragers

**DOI:** 10.1101/2023.11.06.565788

**Authors:** Emilia Moreno, Andrés Arenas

**Affiliations:** Laboratorio de Insectos Sociales, Departamento de Biodiversidad y Biología Experimental, Facultad de Ciencias Exactas y Naturales, Universidad de Buenos Aires, Buenos Aires, Argentina; Instituto de Fisiología, Biología Molecular y Neurociencias (IFIBYNE), CONICET – Universidad de Buenos Aires, Buenos Aires, Argentina

**Author notes:** **SUMMARY STATEMENT**: Honey bees that specialize in foraging for pollen or nectar differ in their perception of sugar and pollen rewards, leading to inter-individual differences in learning that contribute to task specialization.

## Abstract

Social insects live in communities where cooperative actions heavily rely on the individual cognitive abilities of their members. In the honey bee (*Apis mellifera)*, the specialization in nectar or pollen collection is associated with variations in gustatory sensitivity, affecting both associative and non-associative learning. Gustatory sensitivity fluctuates as a function of changes in motivation for the specific floral resource throughout the foraging cycle, yet differences in learning abilities between nectar and pollen foragers at the onset of recollection remains unexplored. Here, we examined nectar and pollen foragers captured upon arrival at food sources. We subjected them to an olfactory PER conditioning using a 10% sucrose solution paired (S10%+P) or unpaired (S10%) with pollen as a co-reinforcement. For non-associative learning, we habituated foragers with a 10% sucrose solution paired (S10%+P) or unpaired (S10%) with pollen, followed by dishabituation tests with either S50% or S10%+P. Our results indicate that pollen foragers show lower performance than nectar foragers when conditioned with S10%. Interestingly, performance improves to levels similar to those of nectar foragers when pollen is included as rewarding stimulus (S10%+P). In non-associative learning, pollen foragers tested with S10%+P displayed a lower degree of habituation than nectar foragers and a higher degree of dishabituation when pollen was used as the dishabituating stimulus (S10%+P). Altogether, our results support the idea that pollen and nectar honey bee foragers differ in their perception of rewards, leading to inter-individual differences in learning that contribute to foraging specialization.

## INTRODUCTION

Social insects inhabit organized societies wherein their cooperative actions depend to a large extent upon the distinctive cognitive abilities of each member (Dukas, 1998; Couzin, 2009; Couzin, et al., 2011; Sih and Del Giudice, 2012). Division of labor is a key feature of social insects, as cohorts of specialized sterile workers engage in diverse tasks simultaneously, enabling colonies to function efficiently. Division of labor is explained through the response threshold model, positing that colony members exhibit variability in their sensitivity (and therefore in their responsiveness) to biologically relevant stimuli associated with specific tasks (Robinson, 1992; Bonabeau et al., 1996). Hence, task allocation of an individual worker is an emergent property of the colony that arises from the interplay between genetic predisposition and epigenetic regulation, ultimately impacting on its abilities to perceive and consequently learn task-related stimuli.

Pollen (protein supply) and nectar (carbohydrates supply) are the main stimuli driving the foraging behavior of the honey bee *Apis mellifera* (Seeley, 1995). Despite flowering plants generally offer both resources at the same time as rewards, honey bee foragers largely specialize in collecting either pollen or nectar (Winston, 1987). Such specialization is probably the best studied case that assesses variations in behavioral responsiveness within the framework of division of labor (Page et al., 2006) and it has been correlated with variations in sensitivity to sucrose, a main constituent of nectar. Furthermore, differential responsiveness or sensitivity to rewards affects learning performance (Scheiner et al., 2001a; 2001b; 2005), and consequently, worker task allocation within the colony.

Honey bees’ sucrose sensitivity can be estimated by the lowest concentration of sucrose solution that triggers the proboscis extension reflex (PER). This innate response occurs when beeś antenna come into contact with an enough concentrated sugar solution (Minnich, 1932; Marshall, 1935; Kuwabara, 1957; Takeda, 1961; Page et al., 1998; Pankiw and Page, 1999). Pioneer studies evaluated the sucrose responsiveness of bees captured returning to the hive. They found that pollen-collecting bees were more responsive across a range of sucrose concentration compared to nectar foragers (Page et al. 1998; Pankiw and Page 1999, 2000; Scheiner et al. 2001b, 2003a). However, a recent study evaluated sucrose sensitivity of pollen and nectar foragers captured immediately after initiating resource recollection or upon arrival at a food source and revealed a reduced responsiveness of pollen foragers compared with nectar foragers at the end of the recollection cycle (Moreno and Arenas, 2023). These findings indicate an interplay between foraging motivation and predisposition to search for nectar or pollen, influencing foragerś gustatory sensitivity.

In addition to innate responses triggered by pollen and nectar (Dobson et al., 1999; Arenas and Farina, 2014; Galizia et al., 2005; Raguso, 2008; Carr et al., 2015), food sources provide other cues, such as odors, colors, shapes, etc., that enhance foraging efficiency after being learned by the bees. In turns, associations between sensory cues and rewards lead to memories that impacts orientation, landing responses (Chaffiol et al., 2005; Nery et al., 2020; Arenas et al., 2007; Arenas and Farina, 2008), and trigger the extension of the proboscis (Takeda, 1961; Bitterman et al., 1983). As a result, foraging behavior of honey bees is underpinned by associative learning processes akin to Pavlovian classical conditioning, where the pairing of a conditioned stimulus (CS), such as floral cues, with an unconditioned stimulus (US), like a nectar reward, established a predictive relationship wherein individuals anticipate the US solely upon the presentation of the CS. In laboratory settings, the extension of the proboscis can be olfactory conditioned through repeated pairings of a neutral odor and a sufficiently concentrated sucrose reward (Minnich, 1932; Marshall, 1935; Kuwabara, 1957; Takeda, 1961; Bitterman et al., 1983).

Considering that the intensity of both the conditioned stimuli (CS) and the unconditioned stimuli (US) is relevant to establish an association (Rescorla and Wagner, 1972; Pelz et al., 1997), the way bees perceive stimuli plays a pivotal role in their learning performance (Moreno et al., 2022). Notably, bees with a heightened sucrose sensitivity learn better during conditioning than less sensitive individuals (Scheiner et al., 2005). Moreover, conditioning performance of pollen foragers rewarded with sucrose, is lower upon arrival at the food source than upon departure (Moreno and Arenas, 2023). Interestingly, such a difference became not detectable when bees were co-reinforced with pollen (Nery et al., 2020; Moreno and Arenas, 2023), suggesting that perception of pollen and nectar is dynamic throughout the foraging visit. In a previous study, Nery and co-workers (2020) compared learning and memory retention of pollen foragers captured when returning to the nest, with that of nectar foragers captured shortly after their arrival at a feeder. In this scenario, where sucrose sensitivity is expected to be more similar between the two groups (Moreno and Arenas, 2023), pollen foragers learned better and extinguished memories less than nectar foragers if conditioned with a combination of sucrose and pollen. Interestingly, both groups displayed similar learning performances when conditioned with sucrose alone (Nery et al., 2020). It is important to note that, until now, differences in learning between nectar and pollen foragers have not been explored with bees captured early in the foraging visit, when they would be highly motivated to forage.

Beyond associative learning, bees also exhibit non-associative learning if they are repeatedly exposed to a meaningful event or stimulus. Such is the case of habituation, where the repeated presentation of an unconditioned stimulus (US) leads to a gradual reduction in the intensity or likelihood of a response (Humphrey’s, 1933; Thompson and Spencer, 1966). Behavioral responses exhibiting habituation may encompass a spectrum of outcomes within the nervous system, ranging from basic reflexes and muscle contraction to the release of hormones or the activity of motor neurons (Rakin et al., 2009). Therefore, this type of learning has an important adaptive value by preventing animals from keeping responding to stimuli that lack true meaning (Rakin et al., 2009). Habituation is distinguished from simple fatigue or sensory adaptation because the response can be restored to its initial levels through the presentation of the same stimuli at higher concentrations, or with novel equivalent stimulation, a process known as dishabituation (Thompson and Spencer, 1966). In honey bees, the repeated presentation of a low-concentrated sucrose solution induces habituation of the PER. Habituation, in turn, can be dishabituated by presenting a higher-concentrated sucrose solution (Scheiner, 2004). Notably, the degree of habituation in honey bees correlates with their sucrose sensitivity: bees highly responsive to sucrose display less habituation and a greater degree of dishabituation compared to bees with lower sucrose responsiveness (Scheiner, 2004). So far, there is no available evidence regarding the role of pollen as a habituating or dishabituating stimulus. Furthermore, no studies have explored potential differences in the degree of habituation and/or dishabituation between pollen and nectar foragers.

Given the perception of nectar and pollen rewards influenced by the stage of the foraging cycle and the forager’s predisposition (Moreno and Arenas, 2023), our study searched for differences in learning abilities that contribute to understanding the division of labor between pollen and nectar foragers. To this end, we investigated both associative and non-associative learning in honey bee foragers captured at the beginning of their foraging visit, a critical moment as the sensory perception of the bee is finely tuned to achieve the collection of either pollen or nectar. In a first experiment, we assessed acquisition performance during the classical olfactory PER conditioning, with and without the presentation of pollen as a co-reinforcement. Following the conditioning phase, we evaluated the retention of olfactory memories 24 hours later by presenting either the learned odor (CS+) or a novel odor (CS-). If forager’s perception depends on its motivation for searching pollen or nectar, we predict that: (i) nectar foragers will exhibit enhanced learning performances and higher memory retention levels when conditioned with a sucrose reward compared with pollen foragers, and (ii) the performance of pollen foragers will improve when pollen participates as an additional rewarding stimulus. In our second experiment, we subjected foragers to PER habituation using various combinations of sucrose concentrations and sucrose + pollen as habituation and dishabituation stimuli. We predict that upon beeś arrival: (i) pollen foragers will display reduce habituation when exposed to sucrose + pollen as habituated stimuli compared to nectar foragers, and (ii) pollen foragers will display a higher degree of dishabituation than nectar foragers when pollen participate as dishabituaing stimulus.

## METHODS

We tested European honey bees *A. mellifera ligustica.* Foragers were trained to visit a foraging station located 150 m away from the apiary, where *ad libitum* feeders with sucrose 10% w/w and crushed bee-collected multifloral pollen were located separately (20 cm apart from each other) on a wooden platform (30cm x 40cm). We captured pollen and nectar foragers upon arrival, immediately after they started ingesting sucrose solution or collecting pollen from the feeders. In the laboratory, we chilled foragers in a refrigerator (−5 °C) and carefully restrained them in metal tubes allowing their first legs and mouthparts to move freely (Kuwabara, 1957; Bitterman et al., 1983). We offered them water until satiation and then placed them in the incubator (33 °C, 60% RH, and darkness) for 20 to 30 minutes.

### Experiment 1: Associative learning

We olfactory conditioned bees by the presentation of the floral odor linalool (0.1 M, Sigma-Aldrich) as conditioned stimulus (CS+) to both antennae, sucrose-water solution (10% w/w) as unconditioned stimuli (US) to both antennae, and hand-collected kiwi pollen as co-reinforcement to the right or to the left tarsus. Bees were olfactory conditioned along 5 acquisition trials (Fig. 1A). Treatments consisted in the presentation of (i) the odor (CS+) paired with sucrose (US) and with pollen (paired procedure, PP; Fig. 1B) or (ii) the odor paired with sucrose but unpaired with pollen (unpaired procedure, UP; Fig. 1C). When the conditioned PER occurred, we assigned values of 1, and when it did not, we assigned values of 0. Bees that showed a spontaneous response (i.e. extending the proboscis in response to the first odor presentation) were excluded, as we cannot determine whether this is an innate response or it indicates a prior (uncontrolled) odor-reward association. Finally, we determined whether forager type and pollen reinforcement affect retrieval of the olfactory memory 24 hours after training. To this end, we tested the conditioned response of trained bees fed with sucrose solution until satiation and maintained 24 hours in the incubator. We presented either the rewarded (linalool; CS+) and a novel, unrewarded, conditioned stimulus (nonanal, CS-) with a 20 min intertrial, both without US (Fig. 1D). We presented the CS+ and CS-in a random order.

**Figure 1.**
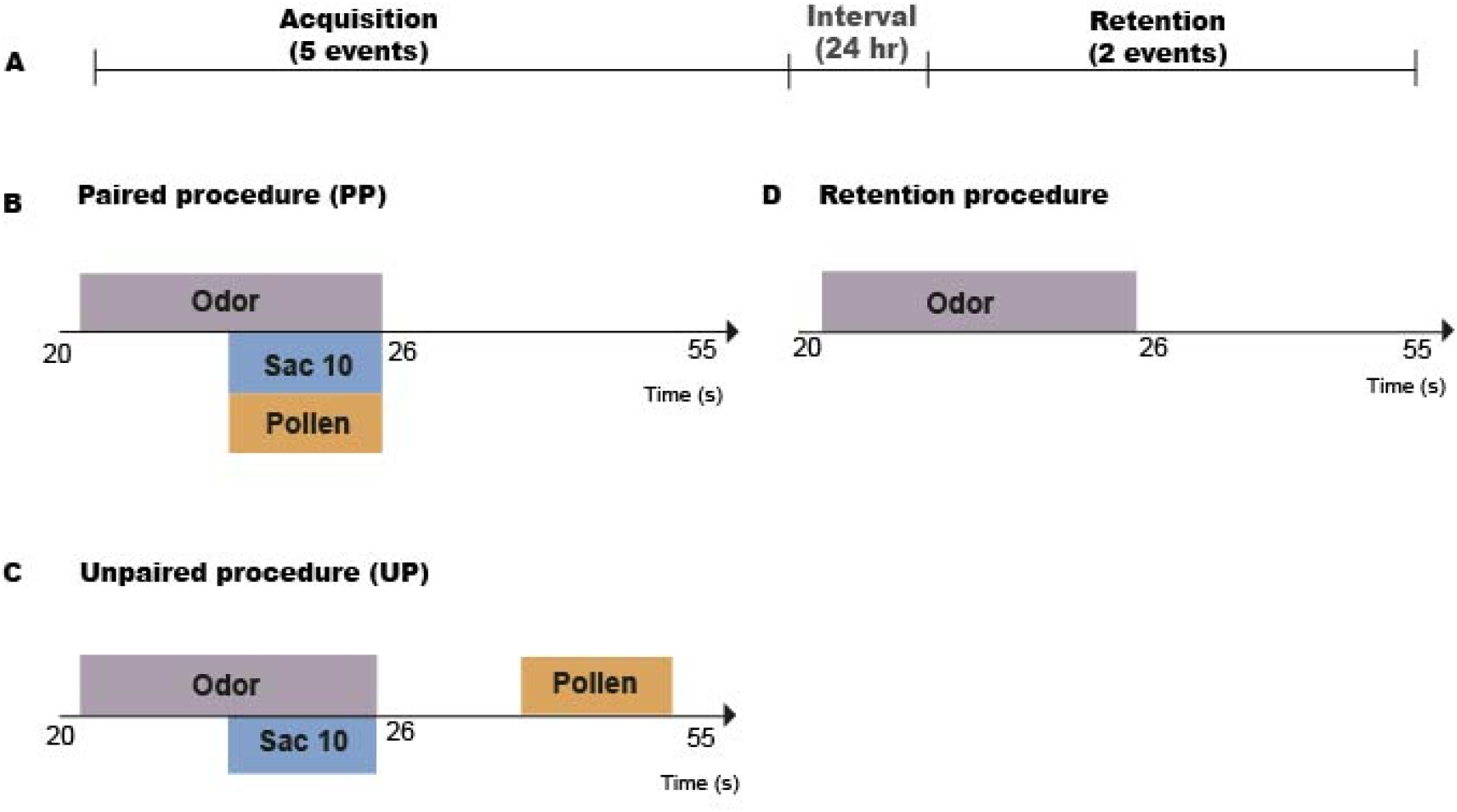
A) Associative learning protocol. Bees were olfactory conditioned to 5 acquisition trials that lasted 55 seconds with an intertrial of 20 minutes. **B) Paired procedure (PP)**. Stimulation consisted in the presentation of the odor on the antenna paired with sucrose solution (10%w/w) on the proboscis and with the contact of a wooden stick wrapped with a cotton layer covered with hand-collected kiwi on the first tarsi. **C) Unpaired procedure (UP)**. The odor was paired with sucrose solution (10%w/w) but unpaired with pollen. **D) Retention procedure.** We tested the retrieval of the olfactory memories 24 hours after training. We presented, randomly, the rewarded odor (linalool; CS+) or the novel odor (nonanal, CS-) with an interval of 20 minutes between each odor presentation. Each trial lasted 55 seconds.

To present odors, we used a device that sent a continuous clean air flow (50 ml s-1) and delivered the odor through a secondary air stream (6.25 mlLs-1) which was injected into the main airflow through a system of valves controlled by computer. A piece of filter paper (30×3 mm) was impregnated with an aliquot of the odor (4 μL) and placed inside a syringe connected to the secondary air stream. Each trial lasted 55s. The valve opening was programmed so that it released clean air during the first 20s, followed by the odor (6s), and a final exposure to clean air for the last 29s. In the paired procedures, the last 3s of the odor presentation overlapped with the sucrose + pollen presentation. During the unpaired procedure, we presented pollen 5s after the odor + sucrose presentation. We measured the PER during the first 3s of the odor presentation.

### Experiment 2: Non associative learning

We habituated the PER response of the bees through a procedure that consisted in the repeated presentation of 25 trials (Fig. 2A) of: (i) sucrose solution (10% w/w) on both antennae paired with the contact of a wooden stick wrapped with a cotton layer covered with hand-collected kiwi pollen on the first tarsi (S10%+P; Fig. 2B) or (ii) sucrose solution (10% w/w) on both antennae (S10%; Fig. 2C) paired with the contact of the tarsi with a wooden stick wrapped with a clean cotton layer as control for mechanical stimulation. The presentation of each trial lasted 1 second and was followed by an inter-trial of 10 s. When the PER occurred, we assigned values of 1, and when it did not, we assigned values of 0. After 25 habituation trials we waited 2 min before starting with 5 dishabituation events (Fig. 2A). Dishabituation tested in bees habituated with sucrose (10% w/w) + pollen consisted in the repeated presentation (5 trials) of either (i) sucrose water solution (50% w/w) + pollen (S50%+P; Fig. 2D) or (ii) sucrose water solution (50% w/w; S50%; Fig. 2E). Dishabituation tested in bees habituated with sucrose water solution (10% w/w) consisted in the presentation of either (i) sucrose water solution (10% w/w) + pollen (S10%+P; Fig. 2F) or (ii) sucrose water solution (50% w/w) (S50%).

**Figure 2.**
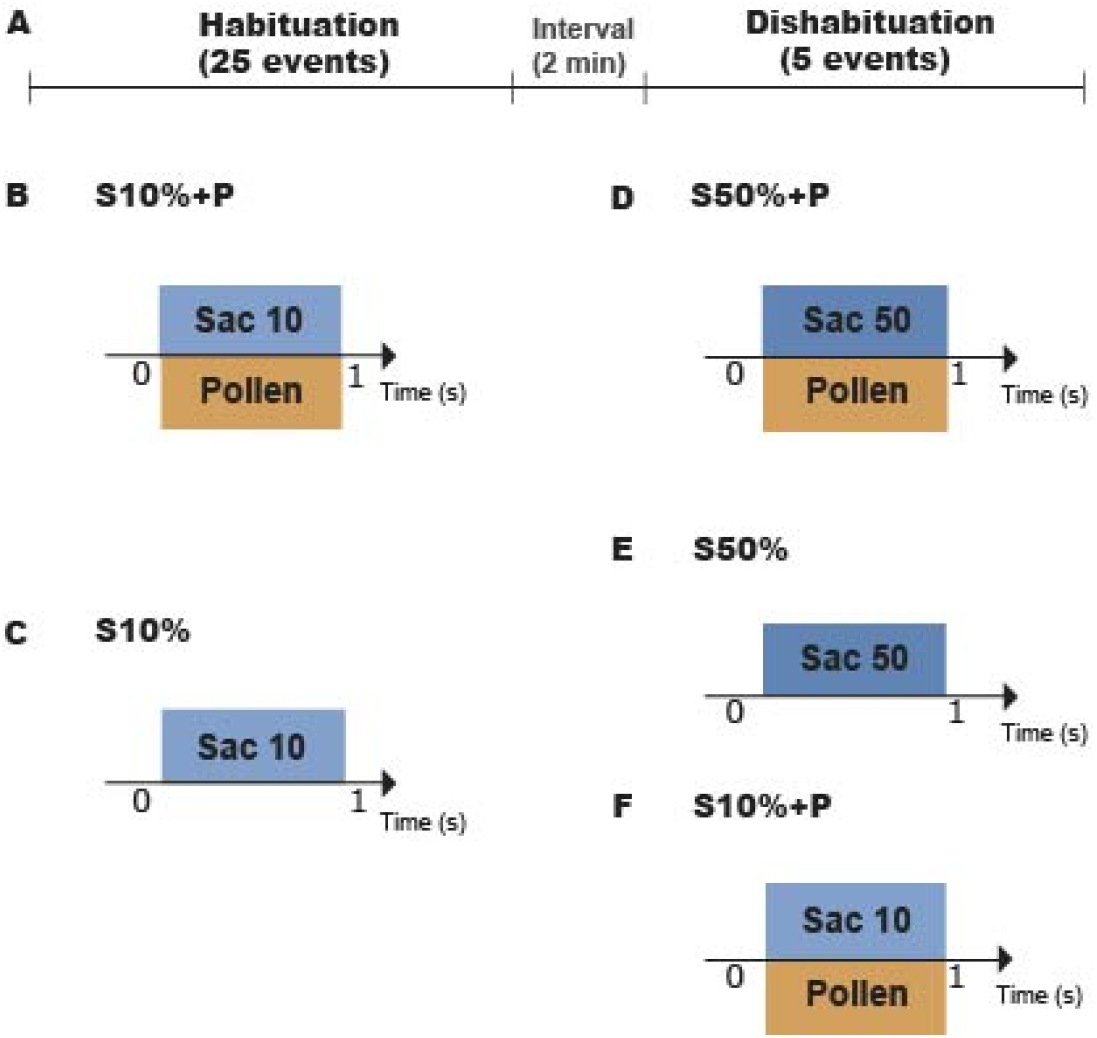
A) Non-associative learning protocol. We habituated the PER response of the bees in 25 trials. After the habituation trials we waited 2 min before starting with 5 dishabituation events. Each stimulation lasted 1 second and was followed by an inter-trial of 10 seconds. **A) Sucrose + pollen procedure (S10%+P).** Stimulation consisted in the presentation of sucrose solution (10% w/w) on both antennae paired with the contact of a wooden stick wrapped with a cotton layer covered with hand-collected kiwi pollen on the first tarsi. **B) Sucrose procedure (S10%).** Stimulation consisted in the presentation of sucrose solution (10% w/w) on both antennae paired with the contact of the tarsi with a wooden stick wrapped with a clean cotton layer as control for mechanical stimulation. **C) Dishabituation procedure of bees habituated with S10%+P**. Dishabituation consisted in the repeated presentation of 5 trials of either sucrose solution (50% w/w) + pollen (S50%+P) or **D)** sucrose solution (50% w/w) alone (S50%). **E) Dishabituation procedure of bees habituated with S10%.** Dishabituation consisted in the presentation of either sucrose solution (10% w/w) + pollen (S10%+P) or sucrose solution (50% w/w) alone (S50%).

### Statistical analysis

All statistical analyses were performed in R (http://www.R-project. org/). In experiment 1, we analyzed the PER proportion of foragers by means of a multiplicative generalized linear mixed model assuming a Bernoulli distribution, using the “glmer” function of the ‘lme4’ package (Bates et al., 2015; Lenth, 2015). For memory acquisition, we considered forager type (a two-level factor corresponding to pollen and nectar foragers), treatment (a two-level factor corresponding to reinforcement procedure: paired pollen and unpaired pollen), and trial (a five-level factor corresponding to each conditioning trial) as fixed effects. Each bee and each day were considered as random effects. For memory retention, we considered forager type (a two-level factor corresponding to pollen and nectar foragers), treatment (a two-level factor corresponding to reinforcement procedure: paired pollen and unpaired pollen) and odor (a two-level factor corresponding to learned and unknown) as fixed effects. Each bee and the day on which experiments were conducted were considered as random effects.

In experiment 2, we analyzed the PER proportion by means of a binomial multiplicative generalized linear mixed model using the “glmmTMB’’ function of the ‘glmmTMB’ package (Bates et al., 2015). We used the proportion of the 25 events together as the response variable. We considered forager type (a two-level factor corresponding to pollen and nectar foragers) and treatment (a two-level factor corresponding to habituation stimulus: S10% and S10%+P) as fixed effects. Each experimental day was considered as random effects. For dishabituation, we used the proportion of the 5 events together as the response variable and the proportion of the habituation treatment as covariable. We analyzed separately bees habituated with S10% from bees habituated with S10%+P. For the first group, we considered forager type (a two-level factor corresponding to pollen and nectar foragers) and dishabituation stimulus (a two-level factor corresponding to S50% and S10%+P) as fixed effects. For the second group (S10%+P), we considered forager type (a two-level factor corresponding to pollen and nectar foragers) and dishabituation stimulus (a two-level factor corresponding to S50% and S50%+P) as fixed effects. Each day was considered as random effects. *Post hoc* contrasts were conducted on models to assess effects and significance between fixed factors using the “emmeans” function of the ‘emmeans’ package version 1.4 (Lenth, 2019) with a significance level of 0.05.

## RESULTS

Our analysis for acquisition detected differences between trials (χ²=59.646, df=13, p=6.076e-08) as well as a double interaction between forager type and reinforcement type (χ²=17.5939, df=4, p=0.001481). Using pollen paired with sucrose as reinforcement (PP), we observed that pollen foragers acquired olfactory memories similar to nectar foragers (z ratio=-1.288, p=0.5707; Fig. 3A). However, when pollen was presented unpaired with the sucrose (UP), we observed that acquisition performance was lower in pollen than in nectar foragers (z ratio=2.654, p=0.0356; Fig. 3A). Acquisition performance of pollen foragers under the paired procedure was higher than under the unpaired procedure (z ratio=4.585, p<.0001). Nevertheless, acquisition performance of nectar foragers under the paired procedure was similar to that under the unpaired procedure (z ratio=1.223, p=0.6123; Fig. 3A). Finally, our analysis for memory retention detected differences in the retrieval performance between the conditioned and the novel odor (χ²= 5.5047, df=1, p=0.01897), but there were no differences related to the forager type nor to the reinforcement type (Fig. 3B).

**Figure 3.**
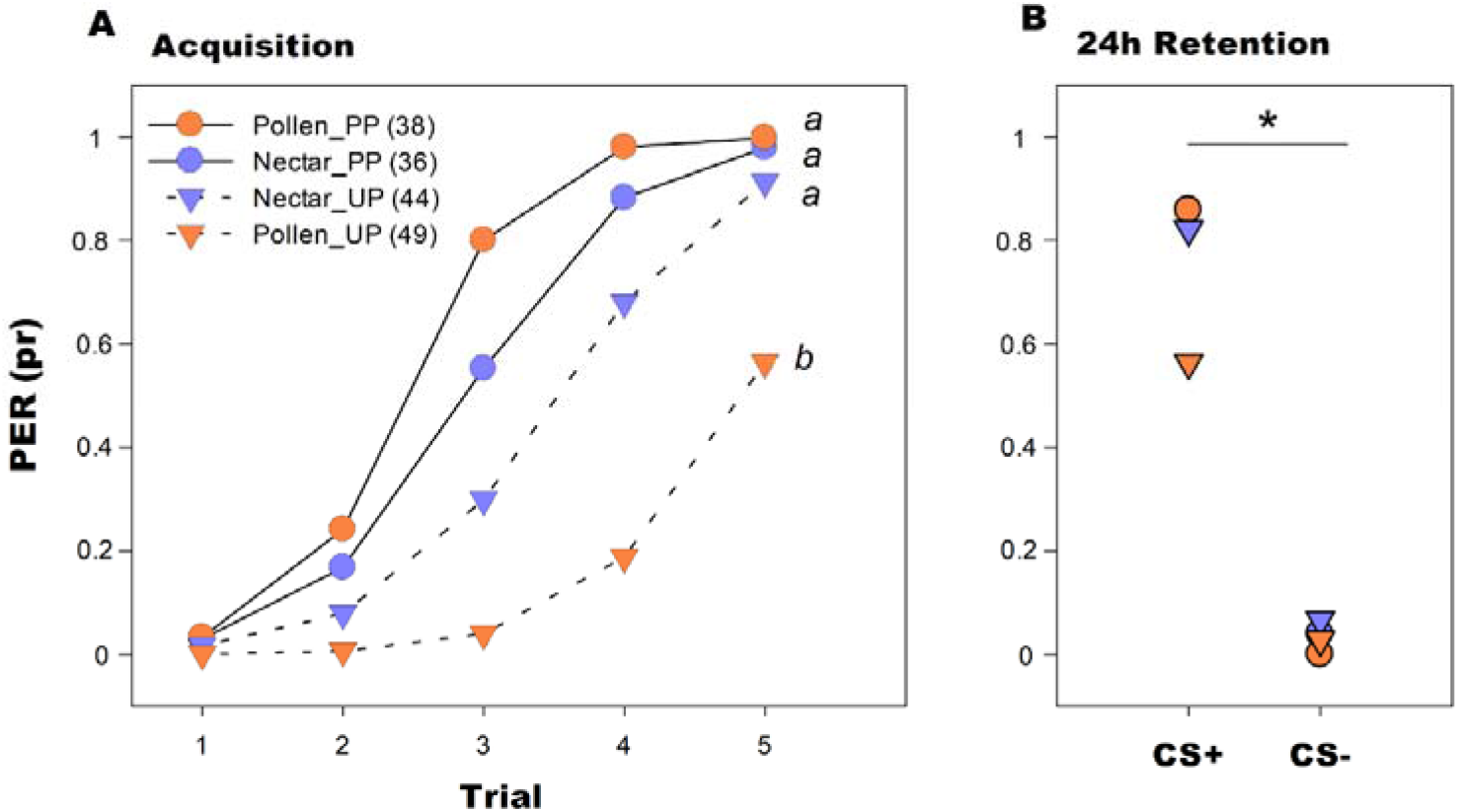
Associative learning. Proportion of PER responses of pollen (orange) and nectar (blue) foragers during **A)** acquisition and **B)** retention trials (predicted data). Pollen was used as co-reinforcement paired (circles) or unpaired (triangles) with the odor (CS+). Different letters indicate statistical differences (p < 0.05). Sample sizes are indicated in parenthesis.

We observed a sustained decline in the PER proportion over the 25 habituating events (Fig. 4A). Habituation occurs faster in bees trained with S10%+P than in those trained with S10%. Consistently, our analysis for habituation detected a double interaction between forager type and reinforcement type (χ²= 6.3244, df=1, p=0.011). When we used S10%+P as habituation stimulus, pollen foragers habituated less (they showed higher PER proportion) than nectar foragers (z ratio=-3.872, p=0.0001), while the PER proportion under the S10% stimulation was similar between pollen and nectar foragers (z ratio=-0.452, p=0.6512; (Fig. 4B).

**Figure 4.**
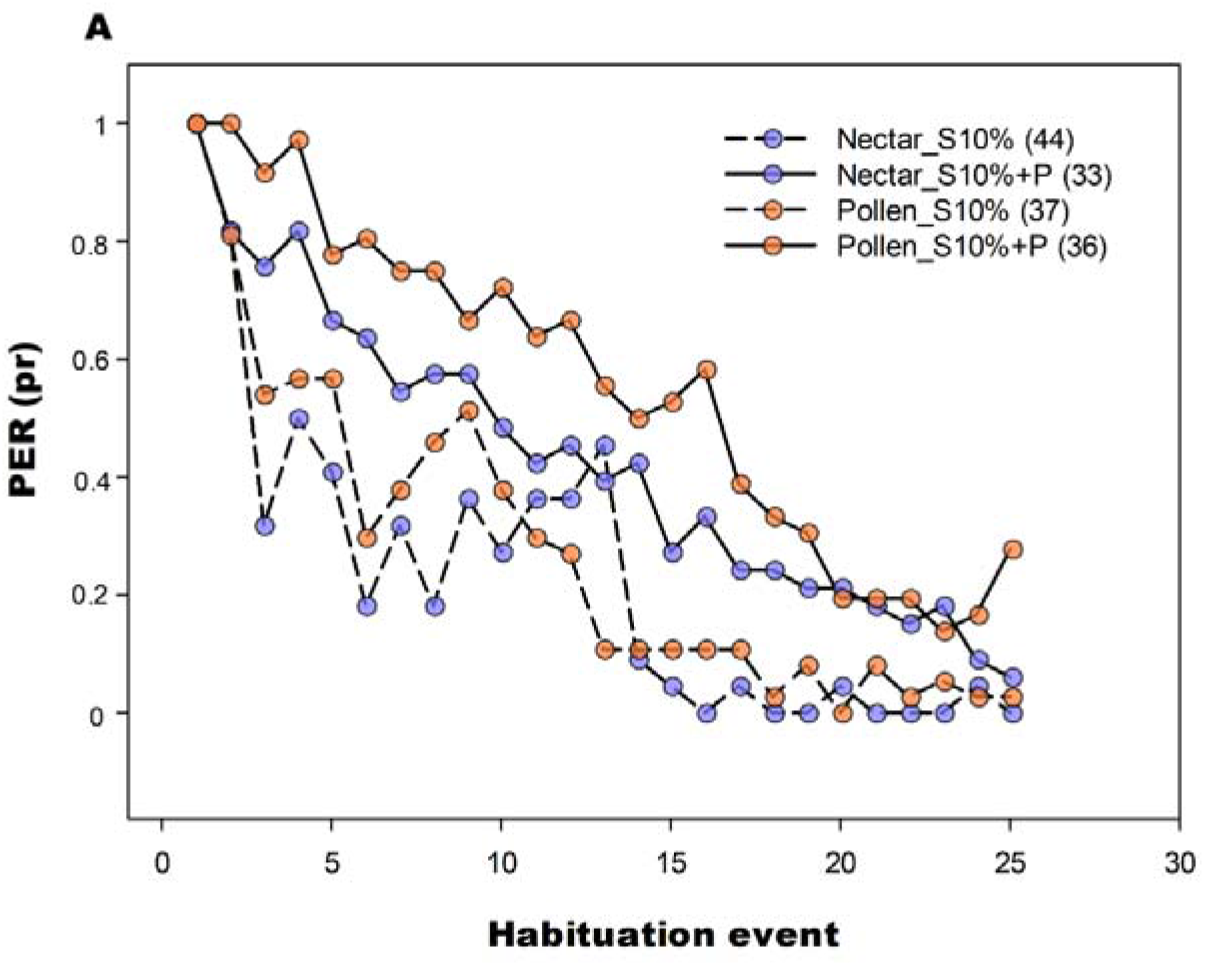
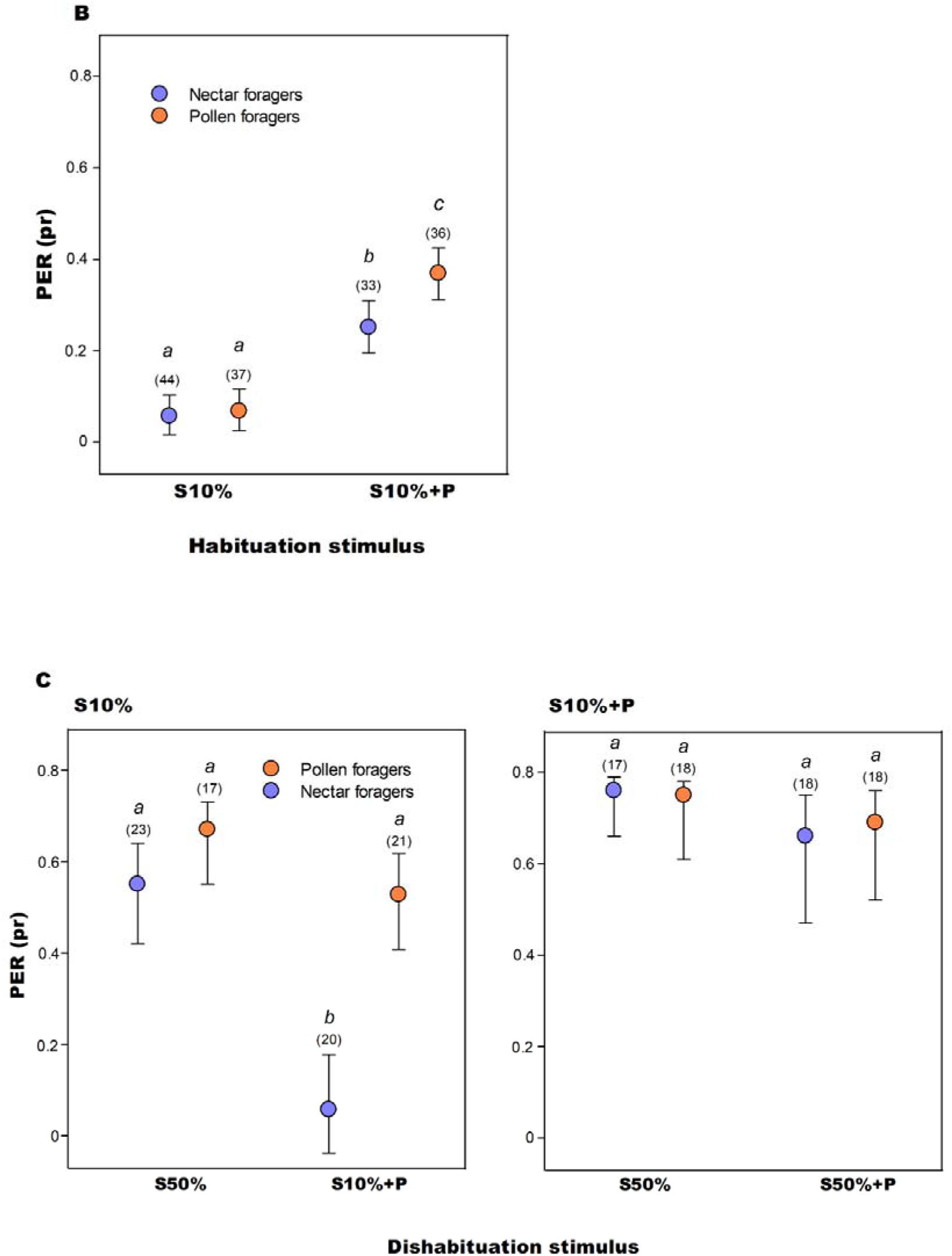
Non-associative learning. **A)** Proportion of PER along 25 habituation trials (observed data) of pollen (orange) and nectar (blue) foragers. We used sucrose or sucrose + pollen as habituation stimuli. **B)** Proportion of PER of all the habituation trials (predicted data) of pollen (orange) and nectar (blue) foragers. **C)** Proportion of PER of 5 dishabituation trials (predicted data) of pollen (orange) and nectar (blue) foragers habituated with S10% (left) and with S10+P (right). Different letters indicate statistical differences (p < 0.05). Sample sizes are indicated in parenthesis.

Dishabituation analysis of bees habituated with S10% detected a double interaction between forager type and dishabituation stimulus (χ²=3.84, df=1, p=0.049; Fig. 4C, left panel). Our analysis detected that the dishabituation performance of pollen and nectar foragers was similar when we used S50% (z ratio=-1.78, p=0.073). Nevertheless, dishabituation performance of pollen foragers was higher than that of nectar foragers when we used S10%+P (z ratio=-5.305, p <.0001), showing the relevance of pollen as excitatory stimulus for pollen foragers.

Dishabituation analysis of bees habituated with S10%+P did not detect differences between forager types (χ²=0.45, df=1, p=0.5012) or between dishabituation stimuli (χ²=0.0003, df=1, p=0.98; Fig. 4C, right panel).

## DISCUSSION

Inter-individual differences in perception and learning are fundamental to the allocation of colony tasks. In this sense, individuals participate in activities mediated by stimuli to which they exhibit heightened sensitivity (Robinson and Page, 1989; Beshers and Fewell, 2001; Perez et al., 2013; Balbuena and Farina, 2020; Mattiacci, 2023). Here, we observed that pollen and nectar foragers show differences in their learning performance according to the type of reward they want. Consistent with the prediction of the response threshold model (Beshers and Fewell, 2001; Robinson, 1992; Bonabeau et al., 1996), pollen foragers that are olfactory conditioned with a sucrose reward exhibited lower acquisition and retention performance than nectar foragers. However, pollen foragers improved their conditioning performances to levels similar to those achieved by nectar foragers, if pollen was presented as a co-reinforcement.

Consistently, during non-associative learning, pollen foragers exhibited diminished habituation and heightened dishabituation compared to nectar foragers if pollen participates as habituating and dishabituating stimuli. This finding supports the notion that pollen elicits stronger responses in bees specialized in pollen collection rather than in nectar foragers (Nery et al., 2020). This differential perception of flower rewarding stimuli might enable the correct acquisition and retention of the environmental cues that predict the presence of the wanted resource (Scheiner et al., 2001a, 2001b; 2005), thereby increasing foraging efficiency. Thus, a tuned perception of the wanted reward affects how individuals “amplify” or “filter” the information that is important (or not) to perform the task efficiently. For example, the repeated presence of pollen on flowers may cause pollen-related cues to lose their meaningfulness for nectar foragers, but not (or to a lesser extent) for pollen-searching bees.

### Variations in perception and learning throughout the foraging cycle

Early experiments exploring the link between foraging specialization and sucrose sensitivity primarily focused on individuals captured at the hive entrance upon their return from foraging expeditions (Page and Fondrk, 1998; Page et al., 2006). These experiments included controlled feeding to minimize differences in sugar satiety among the bees. In such conditions, pollen foragers exhibit a lower sucrose response threshold compared to nectar foragers. This threshold profile was observed not only at the end of the foraging cycle but also from early (Pankiw and Page, 2000) to later stages in the foragers’ lives (Pankiw and Page, 1999). Consistently, foragers from colonies selected over many generations for ‘‘high-pollen-hoarding’’ behavior (Page and Fondrk, 1995) were more responsive to sucrose than bees from colonies selected for ‘‘low-pollen-hoarding’’ behavior (Page et al., 1998; Pankiw and Page, 1999; Scheiner et al., 2001b). Evidence from both selected strains and wild type bees, indicates that sucrose sensitivity has a strong genetic basis (Page et al., 2000; Page and Erber, 2002; Page et al., 2006). Differences in sucrose response thresholds were interpreted as adaptive, arguing that the higher thresholds observed in nectar foragers directed their foraging efforts toward more productive sources, thereby benefiting the colony’s energy gain (Page et al., 1998; Pankiw et al., 2001). However, this interpretation must be appraised in light of our results and a recent study that used bees captured upon arrival at a food source (Moreno and Arenas, 2023). Such a contrasting response pattern indicates that a foragers’ motivation (represented by the phase of the foraging cycle) interacts with the genetic predisposition influencing gustatory sensitivity. These findings underscore the importance of a more detailed exploration of how bees’ responsiveness varies at other stages of the foraging cycle (e.g. immediately before leaving the hive or in the middle of the foraging visit) and even outside it, such as during periods of inactivity or when foragers remain unemployed inside the nest after depletion of the visited food source.

Just like in associative learning, habituation also shows a correlation with sucrose sensitivity, with the most sensitive bees habituating at the lowest rate (Scheiner, 2004).

Consistently with the idea that pollen elicits stronger responses in pollen than nectar foragers (Nery et al., 2020), pollen foragers displayed a lower degree of habituation than nectar foragers and a higher degree of dishabituation if pollen participates as co-reinforcement. However, some results have not been fully in line with what we expected. For instance, habituation to a 10% sucrose solution (S10%) was not lower (but rather similar) in nectar foragers compared to pollen foragers. The probability of response to a 10% sucrose solution was estimated to be around 0.1 for pollen foragers upon arrival at the food source and 0.3 for nectar foragers (Moreno and Arenas, 2023). This difference, depending on data variability, may not be substantial enough to produce an impact during habituation. Conducting further habituation experiments using a broader range of concentrations (including both less and more concentrated solutions) could enhance experimental accuracy to detect differences between forager types. Likewise, the lack of differences in response to the S50% dishabituation stimulus might be attributed to the fact that both forager groups exhibit similar response probabilities (close to 0.85, as observed in Moreno and Arenas, 2023) when confronted with the highly concentrated sucrose solution.

### Variations in perception and learning driven by sugar satiety

The distinctive pattern of sugar response thresholds observed in foragers arriving at the food source and returning to the nest, may be associated with fluctuations in bee sugar satiety throughout the foraging cycle. Foraging is a behavior inherently altruistic, primarily serving the colony’s nutritional requirements rather than individual forager sustenance. In this sense, foragers consume nectar as an energy source prior to departing from the nest, theoretically providing them with sufficient energy to complete the foraging cycle without the need of additional nourishment (Harano et al., 2014). Nectar is ingested at the source and subsequently stored within the digestive tract, specifically the crop, for transportation back to the hive. The collection of nectar satiates foragers, suggesting that besides colony nutritional requirements, the individual forager’s energy demand also plays a role in regulating foraging activity. Consistently, nectar foragers can use the collected nectar to immediately obtain energy if needed (Blatt and Roces, 2001). In contrast, pollen foragers transport pollen on external structures located on their third pairs of legs and use nectar from their crops to bind pollen grains together. Pollen foragers request nectar to their mates before departing from the nest in pursuit of pollen and/or upon returning to the nest following a successful pollen collection (Camazine, 1993). Consequently, the intrinsic nature of pollen and nectar imposes substantial differences in sugar satiety levels between forager types that diverged in opposing directions. Given that bees notably diminished their sensitivity when satiated (Pankiw et al., 2004), the satiety status of bees arriving at a source (low in pollen foragers and high in nectar foragers), could elucidated the observed distinctions in sucrose perception, first, between pollen and nectar foragers (Moreno and Arenas, 2023), and second, among distinct phases of the foraging bout.

Because sucrose sensitivity influences learning performance (Scheiner et al., 2001a; 2001b; 2005; Perez et al., 2013), it becomes possible to explain why foragers arriving at a food source in search for pollen may exhibit poorer performance compared to those seeking for nectar. Following this line of thought, we postulate that sucrose sensitivity of nectar foragers arriving at the food source is similar to that of pollen foragers returning to the hive. Both groups want sugar, either to gather nectar for the colony, or to refuel their energy reserves after pollen collection. Notably, when we evaluated the performance of pollen foragers arriving at the nest with pollen loads in comparison with nectar foragers arriving empty at the food source, no discernible difference was observed during olfactory conditioning rewarded with sucrose (Nery et al., 2020).

### Exploring the role of pollen as a reward

The majority of research focused on understanding perception, learning and memory in honey bees has employed sucrose as reward, with only a limited number of studies investigating the role of pollen as reinforcement (Grüter et al., 2008; Arenas and Farina, 2012, 2014; Nicholls and Hempel de Ibarra, 2013; Nery et al., 2020; Lajad et al., 2021; Moreno and Arenas 2023). Despite little evidence, both pollen and nectar have frequently been treated as equivalent reinforcements. While sugar rewarding pathways operate both pre- and post-ingestion (Sandoz et al., 2002; Nery et al., 202; Muth et al., 2018), the likelihood of pollen acting via ingestion is considerably lower as they are not capable of digesting it (Moritz and Crailsheim, 1987). Our experimental findings demonstrated that pollen presented on the tarsi enhances learning performances in pollen foragers, indicating that it functions as a reinforcement through pre-ingestive chemo-tactile stimulation of the tarsi receptors. To date, the specific pollen components, including proteins, amino acids, lipids and/or fatty acids (Kim and Smith, 2000; Pernal and Currie, 2001; 2002; Arenas and Farina, 2012), responsible of eliciting an excitatory response on bees, as well as the mechanism through which they are processed and encoded in the bee’ brain remain areas of ongoing investigation.

### How do differences in associative and non-associative learning impact foraging specialization?

Pollen-related cues, including their odors, can attract foragers innately (Dobson and Bergstrom, 2000), and/or even after being learned from the pollen-rich diet they consume at early ages (Arenas and Farina, 2014). In light of our results, we expect that nectar foragers become habituated to the presence of pollen-related cues if the source they approach offers only pollen, as may be the case in some plant species like *Papaver*, *Rosa* and *Solanum*, that do not offer nectar (Vogel, 1983). In the case of sources that offer both resources, nectar foragers might benefit from learning pollen cues (as conditioned stimuli) with nectar to assist foraging. Among pollen foragers, loaded corbiculae may represent an important component to reinforce learning and to establish stable and lasting memories. After chemo-tactile stimulation of antennae and tarsi, the absence of pollen on the corbiculae (i.e. no setal displacement; Ford et al., 1981), could demotivate pollen foragers leading to devaluation of pollen as unconditioned stimuli. Then, the resistance of pollen foragers to habituate against chemo-tactile stimulation would be an adaptation to keep pollen foragers responding to cues that predict the resource sought.

### Concluding remarks

Our study provides new insights into the behavioral and physiological processes involved in the control of resource collection in honey bees. The observed differences in learning highlight that nectar and pollen foragers differ in the valuation of the resources offered by flowers. In contrast to prior studies, which showed a positive correlation between sucrose and pollen sensitivity (Page et al., 1998; Scheiner et al., 2004), our results indicate that foragers who are motivated to collect pollen are more sensitive to pollen but less sensitive to sucrose. Thus, pollen contribution as a reward is greater among pollen than nectar foragers.

## ACKNOWLEDGEMENTS

We thank M. J. Corriale for her help with statistical analyses. We also thank Wagner Chaves for their comments and suggestions and Rocio Lajad for her help during the experiments. We also thank the anonymous reviewers for their positive comments and suggestions.

## Data availability statement

All data generated or analysed during this study will be included in this published article [and its supplementary information files] upon acceptance.

